# Storage and transport of labile iron is mediated by lysosomes in axons and dendrites of hippocampal neurons

**DOI:** 10.1101/2024.12.09.627461

**Authors:** Aiyarin Kittilukkana, Asuncion Carmona, Andrea Somogyi, Chalermchai Pilapong, Richard Ortega

## Abstract

Iron dyshomeostasis in neurons, involving iron accumulation and abnormal redox balance, is implicated in neurodegeneration. In particular, labile iron, a highly reactive pool of intracellular iron, plays a prominent role in iron-induced neurological damage. However, the mechanisms governing the detoxification and transport of labile iron within neurons are not fully understood. This study investigates the storage and transport of labile ferrous iron Fe(II) in cultured primary rat hippocampal neurons. Iron distribution was studied using live cell confocal microscopy with a selective labile Fe(II) fluorescent dye, and synchrotron X-ray fluorescence microscopy (SXRF) for total iron distribution. Fluorescent labelling of the axon initial segment and of lysosomes allowed iron distribution to be correlated with these subcellular compartments. The results show that labile Fe(II) is stored in lysosomes within somas, axons and dendrites and that lysosomal labile Fe(II) is transported retrogradely and anterogradely along axons and dendrites. In addition, we have developed a methodological workflow to quantify labile Fe(II) relative to total iron in neurites. This method is based on correlative imaging of fluorescence microscopy of labile Fe(II) combined with quantitative elemental mapping of total iron by SXRF. Quantitative analysis revealed that after Fe(II) exposure, lysosomal Fe(II) accounts for a small but significant percentage of the total iron content in neurites. These result suggest that after exposure to labile Fe(II), iron is mainly present in a non-reactive form in neurons, while the smaller fraction of reactive labile Fe(II) is stored in lysosomes and can be transported along dendrites and axons.

## 1. Introduction

Iron is an essential element required for a wide range of physiological cellular processes including mitochondrial respiration, DNA synthesis, cell signaling, and growth regulation. In the nervous system, iron is essential for neuronal function supporting energy metabolism, neurotransmitter synthesis, myelination, and synaptic activity. However, disruption of iron homeostasis in the brain leading to iron accumulation and abnormal redox balance has been linked to senescence and pathological conditions such as Alzheimer’s and Parkinson’s diseases [1–5]. Iron accumulation is a prominent factor in oxidative stress and protein aggregation in neurodegenerative diseases [1,4,6]. In particular, iron accumulation is observed in the hippocampus of Alzheimer’s disease (AD) patients [7], a brain region early and severely affected in AD.

To protect cells from iron toxicity, iron is stored in the cytosol in a non-toxic form mainly as ferritin. In addition, a kinetically labile iron pool is present in the cytosol at low concentration to rapidly provide iron for iron-dependent cellular functions [1]. Labile iron refers to a reservoir of exchangeable iron found in cells that is coordinated by low molecular weight molecules [8–9]. Labile iron concentration is tightly regulated in cells to balance iron uptake versus deposition into ferritin. However, labile iron levels may increase aberrantly in some pathological conditions. This form of unshielded reactive iron can catalyze redox reactions, such as the Fenton reaction, generating reactive oxygen species (ROS) [10].

Therefore, maintaining a proper regulation of labile iron is essential to minimize its potential contribution to neurodegeneration and aging-related changes [2,9,10]. However, the mechanisms by which labile iron is detoxified in neurons are not fully understood. In particular, whether and how labile iron is transported along neuronal processes such as axons and dendrites remains to be investigated [3,11]. Dendritic and axonal labile iron transport is necessary to avoid iron accumulation in areas where it could lead to oxidative stress and cell damage. Understanding the underlying mechanisms of labile iron transport and its consequences for neurodegenerative diseases is essential for the development of therapeutic approaches targeting iron dyshomeostasis.

In a previous study, we reported the movement of labile iron in neurites using the FerroFarRed™ fluorescent probe in confocal and super-resolution imaging modes [11]. In this study, we aimed to determine how labile iron is stored and distributed along dendrites and axons of hippocampal neurons. In addition, we present the proof-of-principle of a new methodological workflow to assess labile Fe(II) versus total iron content in subcellular compartments using a combination of advanced iron imaging techniques.

## 2. Materials and Methods

### 2.1. Reagents

The neuronal culture included Gibco^TM^ Neurobasal^TM^ medium with B-27TM Plus Supplement (NB-B27+) from Thermo Fisher Scientific and BrainPhys^TM^ neuronal culture medium from STEMCELL Technologies. Neurobasal^TM^ medium with B-27TM Plus Supplement was used to maintain standard primary rat neuronal cultures until iron exposure, which was performed using BrainPhys^TM^ neuronal culture medium, a serum-free basal medium that mimics physiological conditions in the brain to establish reproducible exposure conditions. Tyrode’s solution was prepared using ultra trace elemental analysis grade water (Fisher Chemical) and was used for cell washing during the exposure steps and for confocal microscopy observations. Tyrode’s solution was formulated using the following ingredients: D-glucose (1 mM), NaCl (135 mM), KCl (5 mM), MgCl_2_·6H_2_O (0.4 mM), CaCl_2_·6H_2_O (1.8 mM), and HEPES (20 mM), all chemicals were purchased from Sigma Aldrich. A solution of ammonium acetate (Sigma Aldrich) was prepared with ultra trace elemental analysis grade water (Fisher Chemical) and used to wash out extracellular salts prior to cryofixation for SXRF experiments. The osmolarity of Tyrode’s and ammonium acetate solutions was adjusted to 235 mOsm, corresponding to the osmolarity of the BrainPhys^TM^ medium. Osmolarity measurements were performed using a vapor pressure osmometer (VAPRO 5600, Elite). The pH of both solutions was adjusted to 7.4, which corresponds the pH of the culture medium. For Fe(II) treatment, neurons were exposed to ferrous ammonium sulfate hexahydrate (FAS) from Sigma Aldrich. BioTracker Far-red Labile Fe(II) Dye (Sigma Aldrich), also known as FerroFarRed^TM^ from Goryo Chemical Inc, was used to detect labile Fe(II) in living neurons with spectral characteristics of maximum absorbance at 646 nm and maximum emission at 662 nm. LysoTracker^TM^ Red DND-99 (Thermo Fisher) was used to label lysosomes *in vitro* with maximum spectral excitation at 577 nm and emission at 590 nm. Red Extracellular Anti-Pan-Neurofascin antibody FL490 conjugate (A12/18) (AntibodiesInc) was used to label axons and to perform live-cell confocal microscopy with spectral maximum excitation at 491 nm and emission at 515 nm.

### 2.2. Cell cultures

For confocal microscopy experiments, cell cultures were performed according to previously published protocols [11]. Briefly, primary rat hippocampal neurons were cultured on glass coverslips and grown in Petri dishes containing an astrocyte feeder layer with NB-B27+ medium until reaching DIV7 (days in vitro 7), which provides a suitable environment for neurons to mature and to develop neurite outgrowths [12]. Neurons were maintained in culture at 37°C in a 5% CO_2_ atmosphere until observation.

For SXRF imaging experiments, primary rat hippocampal neurons were cultured directly on silicon nitride membranes of 500 nm thickness (Silson, UK) in Petri dishes containing an astrocyte feeder layer with NB-B27+ medium until DIV7 according to previously published protocols [13]. Neurons were maintained in culture at 37°C in a 5% CO_2_ atmosphere.

### 2.3. Confocal microscopy

At DIV7, primary rat hippocampal neurons were exposed to FAS in BrainPhys^TM^ culture medium at a final concentration of 100 µM for 1.5 hours at 37°C. After Fe(II) exposure, the cells were washed with Tyrode’s solution to remove any excess iron. Then Fe(II) staining was performed using 5 µM of Ferro-FarRed^TM^ in BrainPhys^TM^ culture medium during 1h at 37°C. For dual color fluorescence imaging of labile Fe(II) and lysosomes, 30 minutes after Ferro-FarRed^TM^ initial exposure, neurons were co-exposed to 75 nM LysoTracker^TM^ during 30 minutes. The LysoTracker^TM^ fluorescence images were used for lysosome counting and lysosome size determination without or with 1.5 h FAS exposure. For dual color fluorescence imaging of labile Fe(II) and axons, 45 minutes after Ferro-FarRed^TM^ initial exposure neurons were co-exposed to 1:150 µL dilution of Anti-Pan-Neurofascin antibody during 15 minutes. After labeling, the coverslips were washed with pre-warmed Tyrode’s solution to remove any traces of reagents that might interfere with the observations. The samples were mounted in a DMI6000 TCS SP8 X confocal microscope (Leica) with temperature control (37°C), motorized stage and HC PL APO 63x/1.40 OIL CS2 objective (Leica) for live confocal imaging in static and dynamic modes (z-stack or time-lapse sequences).

### 2.4. Synchrotron X-ray fluorescence (SXRF) imaging

For correlative imaging of Fe(II) and total iron, primary rat hippocampal neurons were cultured on silicon nitride membranes and exposed to 100 µM FAS in BrainPhys™ for 1.5 hours. Following Fe(II) exposure, labile Fe(II) was labeled using 5 µM FerroFarRed^TM^ during 1 h and fluorescence imaging was observed by confocal microscopy after washing cells with pre-warmed Tyrode’s solution. The silicon nitride membranes were then rinsed with ammonium acetate solution to remove extracellular salts, cryofixed using an automated vitrification system, Vitrobot Marck IV apparatus from FEI (USA), and freeze-dried using a Christ alpha 2-4 LD plus freeze dryer, according to previously published protocols [13]. After freeze drying, FerroFarRed^TM^ fluorescence was observed by epifluorescence microscopy to confirm the location of the regions of interest (ROIs) of Fe(II) puncta initially observed by confocal imaging on living neurons.

SXRF imaging was performed at the NANOSCOPIUM beamline [14], of the French national facility synchrotron SOLEIL. SXRF element distributions were acquired by using a 12 keV X-ray beam focused to 250 x 250 nm^2^ size with a Kirkpatrick-Baez nano-focusing mirror-pair. The continuous FLYSCAN technique has been used for data collection [15]. For each neuron of interest, SXRF imaging was performed with a pixel size of 250 nm and an integration time of 100 milliseconds per pixel. A home-made MATLAB code was used for on-line data treatment at the beamline for extracting the elemental distribution maps from the identified X-ray region of interests.

### 2.5 Quantification of Fe(II) over total iron content

SXRF images of total iron distribution were compared with those of Fe(II) localization obtained by FerroFarRed™ fluorescence microscopy after freeze-drying. In the SXRF total iron maps, a ROI encompassing entire dendrites were selected. Then within this ROI of the SXRF total iron map, smaller ROIs were selected encompassing only the Fe(II) puncta identified thanks to the Fe(II) FerroFarRed^TM^ fluorescence images superposition. SXRF total iron and Fe(II) fluorescence image superposition was performed with ICY software (http://icy.bioimageanalysis.org) and eC-CLEM plugin [16]. The total iron intensity in the ROI encompassing the dendrite and the total iron intensities in the smaller ROIs corresponding to Fe(II) locations were then measured using the FIJI program. The ratio of the sum of the iron intensity in the ROIs corresponding to Fe(II) to the total iron intensity in the whole dendrite was then calculated, providing critical insight into the quantitative distribution of Fe(II) within neurons.

## 3. Results and Discussion

### 3.1. Co-localization of labile iron and lysosomes throughout soma and neuronal processes

We performed confocal imaging of lysosomes and of labile iron in living neurons at DIV7 (Figure 1). FerroFarRed™ is a highly selective Fe(II) turn-on fluorescent probes developed for intracellular localization of labile iron that has been successfully used to detect Fe(II) species within cellular organelles [17–18]. The results of correlative confocal imaging showed that labile iron (red) is almost fully co-localized with lysosomes (green) throughout the soma and neuronal processes in primary rat hippocampal neurons exposed to 100 µM FAS for 1.5 hours (Figure 1).

**Figure 1.**
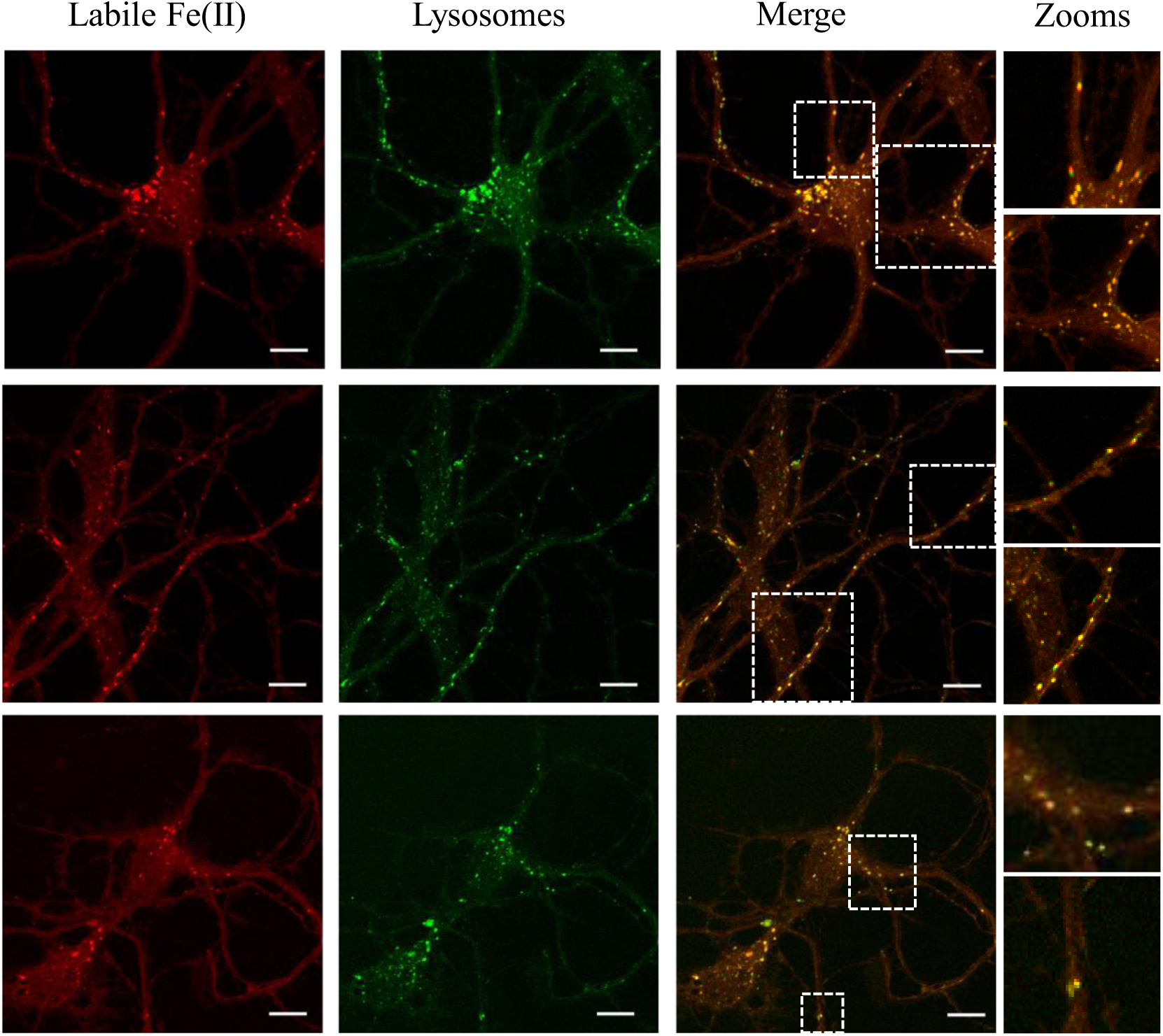
Labile Fe(II) is co-localized with lysosomes in primary rat hippocampal neurons exposed to ferrous ammonium sulfate. Labile Fe(II) is co-localized with lysosomes in the soma and neuronal processes of primary rat hippocampal neurons, as indicated by the predominance of yellow puncta. In some cases, the red and green signals are slightly shifted in the merged image due to the delay between the two acquisitions at two excitation wavelengths, indicating the rapid movement of some of these structures (see zoomed areas). Some lysosomes appear only in green, indicating that not all lysosomes contain Fe(II) (see zoomed areas). Scale bars = 10 µm.

Lysosomes are cellular organelles with an acidic lumenal pH (4.5 to 5) that contain hydrolytic enzymes responsible for the degradation and recycling of cellular waste. Lysosomes play a key role in iron homeostasis by degrading iron-containing proteins such as ferritin or transferrin, which contributes to the tightly regulated release of labile iron in the cytosol [19–21]. Lysosomes receive iron-proteins from the endocytic and autophagic pathways [21]. The acidic environment within lysosomes facilitates the release of iron from ferritin and other macromolecules, making it available for cellular functions. Lysosomes play a key role in converting ferric iron Fe(III) into ferrous iron Fe(II), which can be used by the cell [21–22]. This conversion is essential as many cellular functions, such as mitochondrial respiration and DNA synthesis require Fe(II). In neurons, lysosome-mediated degradation of ferritin is essential to maintain adequate levels of labile iron Fe(II) in the cytosol to meet the high iron demands of neuronal function and neurotransmission. The transporter solute carrier family 11 member 2 (SLC11A2), also named divalent metal transporter 1 protein (DMT1), transports Fe(II) from the lysosome to the cytosol [20–21].

Our results indicate that lysosomes are the major organelle of labile Fe(II) storage in neurons. They also suggest that in addition to DMT1 transport of Fe(II) from lysosomes to the cytosol, Fe(II) is also transported from the cytosol to the lysosome when cells are exposed to extracellular Fe(II). This result highlights that mechanisms other than endocytosis and autophagy may be involved in iron uptake from the cytosol into the lysosomal lumen in the presence of excess Fe(II). Our results also suggest that Fe(II) is rapidly transported from the extracellular space within neurons. Previous studies have shown that primary rat hippocampal neurons exhibit significant iron uptake from transferrin-bound iron (TBI), but also from non-transferrin-bound iron (NTBI), either as Fe(II) or Fe(III), with Fe(II)-NTBI being the most efficient uptake pathway [23]. This intracellular uptake is driven by ZIP8, a Fe(II) iron transporter [23], which is highly expressed in the plasma membrane of both the soma and neuronal processes. Our results showing intracellular Fe(II) uptake and spread out distribution are consistent with a rapid transport of NTBI Fe(II) into the soma and along neuronal processes.

### 3.2. Effect of Fe(II) exposure on the number and size of lysosomes

Next, we sought to determine whether Fe(II) exposure could induce lysosomal changes in primary rat hippocampal neurons. Using LysoTracker^TM^ fluorescence confocal imaging, we measured the mean diameter of lysosomes in dendrites and somas. The results showed that the size of lysosomes was unchanged in response to FAS exposure (100 µM, 1.5 h) (Figure 2a). We also quantified the number of lysosomes per surface area (µm²) within somas and dendrites, with and without FAS exposure, and showed no significant change in this number (Figure 2b).

**Figure 2.**
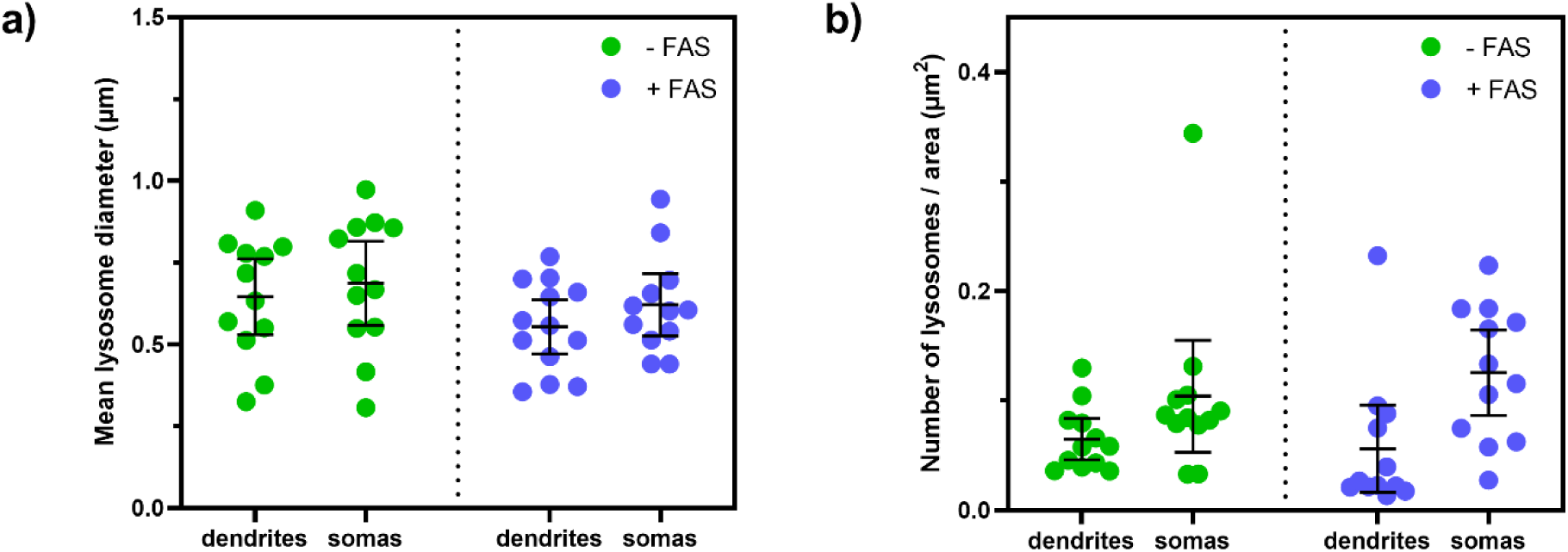
Mean diameter and number of lysosomes in primary rat hippocampal neurons with or without Fe(II) exposure. a) Mean lysosome diameter (in µm) in dendrites and somas of neurons without (-FAS) or with (+FAS) exposure to 100 µM FAS for 1.5 h. Each point represents the mean lysosome diameter of all lysosomes from dendrites and somas of single neurons (n=12 neurons). No significant differences between groups. b) Number of lysosomes per unit area (µm^2^) in somas and dendrites of neurons without (-FAS) or with (+FAS) exposure to 100 µM FAS for 1.5 h. Each point represents the number of lysosomes per unit area counted in dendrites and somas of single neurons (n=12 neurons). No significant differences between groups.

Neurons are particularly susceptible to iron-induced oxidative stress and lysosomes play a protective role by sequestering excess iron [21]. In lysosomes the labile iron pool is buffered via the formation of chelate complexes with low molecular weight molecules such as glutathione and cysteine able to bind Fe(II). Labile iron can also be complexed to polyamines in the lysosomes [21]. In case of chronic Fe(II) exposure the iron-buffering capacity of the lysosomes maybe overwhelmed resulting in iron-induced oxidative stress and cell damage. Therefore, the rapid lysosomal storage capability of neurons is essential to cope with excess exposure to labile iron. We have shown that Fe(II) is rapidly stored in lysosomes without modification of the number and size of lysosomes suggesting a rapid adaptation of neurons to Fe(II) exposure by mobilizing the existing lysosomal pool. Moreover, labile Fe(II) seems to be mobile along neuronal processes as suggested by the results shown in Figure 1.

### 3.3. Lysosomal labile iron shows bidirectional transport in both dendrites and axons

Our next objective was to determine if lysosomal labile iron is quiescent or transported along neuronal processes, if this transport occurs both in axons and dendrites, and if it is anterograde or retrograde. Each neuron exhibits one axon and multiple dendrites. We used anti-pan Neurofascin to label axons and FerroFarRed™ to label labile iron (Figure 3). Anti-pan Neurofascin antibody targets the extracellular domain of neurofascin and provides an accurate live cell labeling of the axon initial segment [24]. Our results demonstrated that labile iron is transported along the axons either anterogradely, moving away from the cell body (Figure 3a), or retrogradely, moving towards the cell body (Figure 3b). Additionally, labile iron is also transported along the dendrites, as indicated by the absence of anti-pan Neurofascin labeling, in both anterograde (Figure 3c) and retrograde (Figure 3d) directions. Furthermore, we observed that most of the labile iron remains stationary within the axon and dendrites, indicating temporary storage (Figure 3e).

**Figure 3.**
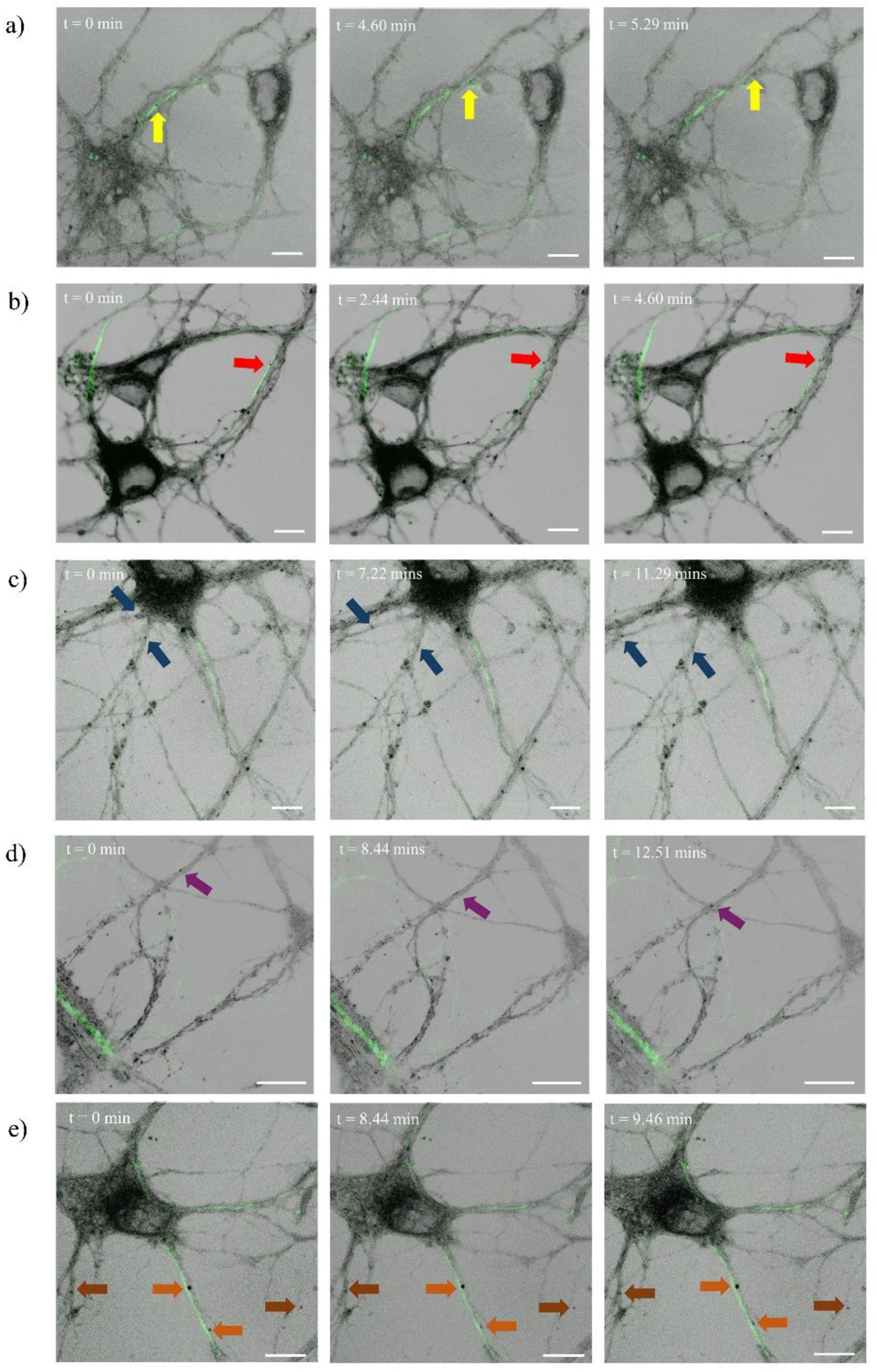
Dual color confocal microscopy of labile iron (black puncta) and axons (green). a) Labile iron transported anterogradely along the axon (yellow arrows). b) Labile iron transported retrogradely along the axon (red arrows). c) Labile iron moved anterogradely along the dendrites (blue arrows). d) Labile iron moved retrogradely along the dendrites (purple arrows). e) Labile iron remains stationary within the axon (orange arrows) and dendrites (brown arrows). Scale bars = 10 µm.

Only a few studies have investigated the dynamic transport of iron along neuronal processes. Anterograde axonal transport of transferrin by retinal ganglion cells from the retina into the optic nerve and optic tract has been observed in animal models using ^59^Fe-tranferrin autoradiography [25]. Another important breakthrough in understanding iron transport in the brain has been provided by the evidence of iron anterograde translocation along specific axonal projections, in particular from ventral hippocampus to medial prefontal cortex to substantia nigra, and from thalamus to amygdala to medial prefontal cortex [26]. These results were obtained on brains from mice using a combination of iron histochemistry with Perl’s staining and ^57^Fe mass spectrometry imaging. Our findings also indicate that labile iron can be transported anterogradely along axons. All together, these results suggest that anterograde axonal transport of transferrin-bound and labile iron could supply iron to distant regions of the brain where iron is needed for neuronal functions. In addition, we provide evidence that anterograde labile iron transport also occurs in dendrites suggesting that iron can be delivered to distal regions of the dendrites within lysosomes. Neuronal function depends significantly on anterograde transport, which ensures that synaptic terminals gain essential components to sustain synaptic transmission and plasticity. Since lysosomal iron is the principal source of iron for many metabolism pathways, in particular for energy production in mitochondria [21], anterograde transport of labile iron might be an efficient source of iron for mitochondria in distal compartments such as synaptic compartments.

We have also observed the axonal and dendritic retrograde transport of lysosomal labile iron. Retrograde iron transport is crucial for maintaining neuronal function in the soma such as mitochondrial energy production. Neuronal somas have a high concentration of mitochondria that require iron as an essential component of several mitochondrial enzymes involved in ATP synthesis. Iron is also essential in the soma for the synthesis of iron-dependent enzymes and proteins. The soma also contains high amount of ferritin that stores iron safely. By transporting iron to the soma, neurons can control iron levels, reducing the risk of iron-induced oxidative stress.

### 3.4. Correlative imaging of labile Fe(II) and total iron

Imaging both labile and total (ferritin-bound) iron in subcellular compartments is usually difficult to perform because it relies on different spectro-imaging methods. We developed a new methodological workflow to compare the intracellular distribution of labile Fe(II) to total iron distribution. This workflow is based on the correlative imaging of labile iron using FerroFarRed™ and epifluorescence microscopy, together with synchrotron X-ray fluorescence imaging of total iron. Representative examples of correlative imaging of total iron and labile Fe(II) distributions are presented in Figure 4, with total iron displayed in yellow and labile Fe(II) in red in the overlay images.

**Figure 4.**
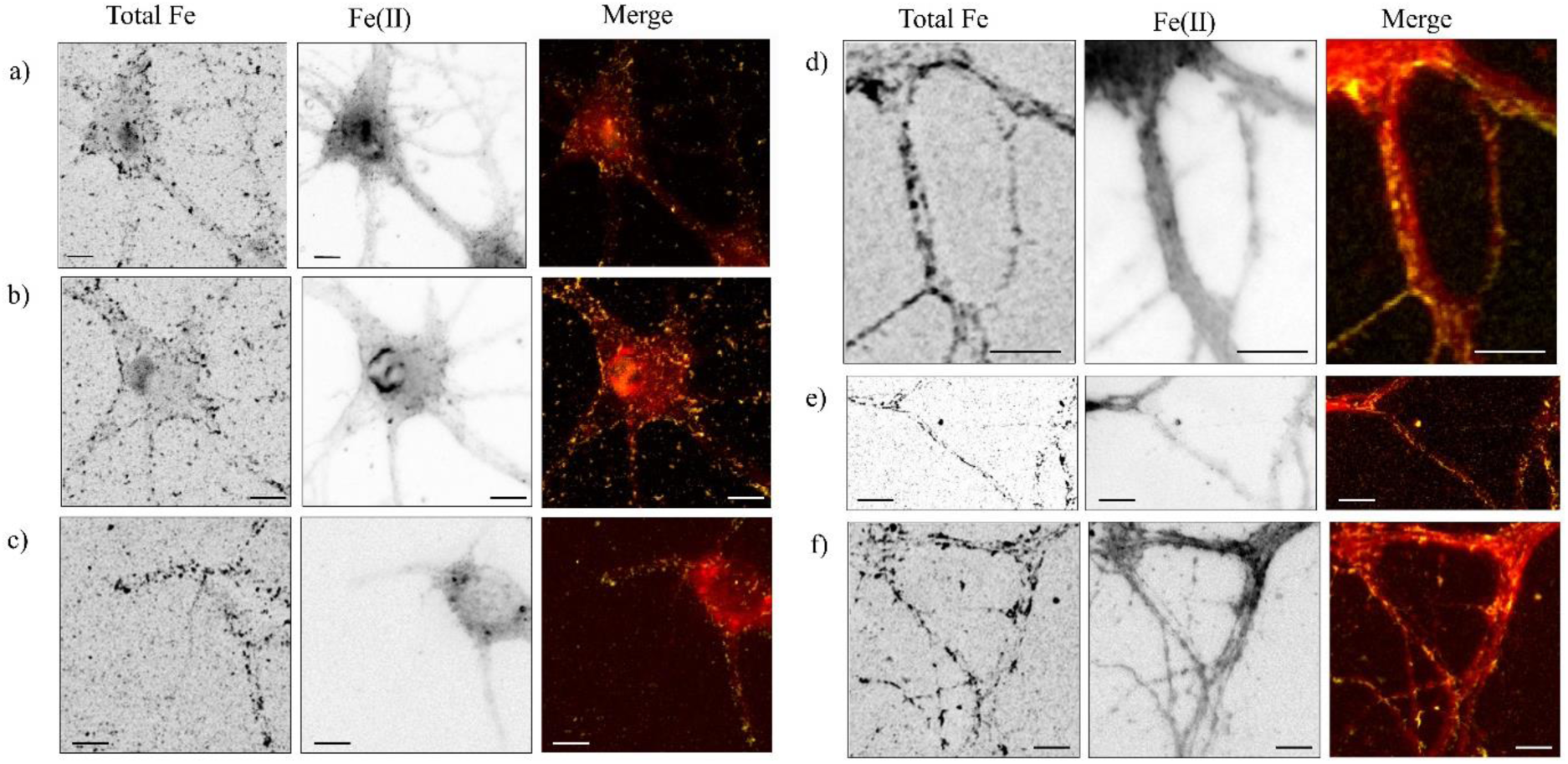
Distributions of total Fe, obtained by SXRF, and of labile Fe(II) using fluorescence of FerroFarRed™, in primary rat neurons exposed to 100 µM FAS, and merged distributions (total iron in yellow and labile Fe(II) in red). Total iron is mainly distributed in form of numerous puncta at the periphery of the soma and along neuronal processes, while Fe(II) puncta are more limited in number and distributed around the nucleus and within some punctual localizations in the cytosol and along neurites. When axons and dendrites are observed more specifically (d-f), Fe(II) puncta are lower in number that total iron structures but are co-localized with these ones as illustrated by the merged images. Scale bars = 10 µm.

The distribution of total iron obtained by SXRF imaging indicates that iron is present in form of micrometric puncta at the periphery of the soma and extends along the neuronal processes (Figure 4, Total Fe). This result agrees with previously published results on total iron distribution obtained by SXRF in hippocampal neurons, showing a dotted distribution of iron in micron scale structures in the cytosol and within dendrites [27–29]. Iron presence throughout neuronal structures is consistent with the established importance of iron in numerous cell functions, such as co-factor of numerous enzymes for neurotransmitter production and mitochondrial energy production. In addition, we found that a large number of iron structures are located at the periphery of the soma after exposure to extracellular labile iron. This result suggests that extracellular iron is rapidly incorporated and stored in structures close to the plasma membrane.

In contrast to the total iron distribution, labile Fe(II) appears less widespread, predominantly localized in the soma as broad signal, and with some localized puncta along the dendrites and axons (Figure 4, Fe(II)). When comparing total iron and labile Fe(II) distributions in neuronal processes (Figure 4 d-f), it appears that labile Fe(II) structures are correlated with total iron structures but are present in lower amounts than these late. Since Fe(II) structures are confined within lysosomes and the majority of lysosomes contained Fe(II) (Figure 1), it can be inferred that most total iron dotted structures are not located in lysosomes and most probably correspond to Fe(III) rich aggregates such as ferritin bound iron. This result suggests that lysosomal incorporation of labile Fe(II) is not the unique pathway developed to protect neurons from Fe(II) toxicity, indicating a rapid conversion to Fe(III) to prevent oxidative stress and maintain cellular integrity. It can also be seen that the strong and broad, non-dotted, FerroFarRed^TM^ signal measured around the nucleus (Figure 4a and 4b), as also observed in live cell imaging (Figure 3), does not correspond to the total Fe distribution but rather to unspecific FerroFarRed^TM^ signal.

### 3.5. Quantification of labile Fe(II) over total iron content

To further investigate the balance of labile iron over total iron content in neurons we performed a quantitative determination based on our correlative imaging method (Figure 5). The ROIs that encompassed the areas of Fe(II) localization in the FerroFarRed™ fluorescence images were selected in the corresponding total iron SXRF images (Figure 5, red circles). Then, the sum intensity of the total iron signal was determined in these ROIs corresponding to the specific Fe(II) puncta. Subsequently, the total iron intensity of the selected dendritic area is measured. By comparing these measurements, the ratio of Fe(II) to total iron can be determined. SXRF images confirm the presence of iron in the Fe(II) puncta observed in the dendrites, as expected from the reactivity of FerroFarRed^TM^. The Fe(II) over total iron fraction results varied depending on the number of Fe(II) puncta and total Fe content observed in the selected dendrites. Despite this local disparities, this method enables to estimate the proportion of Fe(II) in neurons exposed for 1.5 h to 100 µM FAS, which is in the 2.4% to 16% range. In neurons not exposed to FAS, the number of Fe(II) puncta detected is very low, one or two puncta per entire neuron [11], indicating that in control conditions the labile Fe(II) pool is almost undetectable.

**Figure 5.**
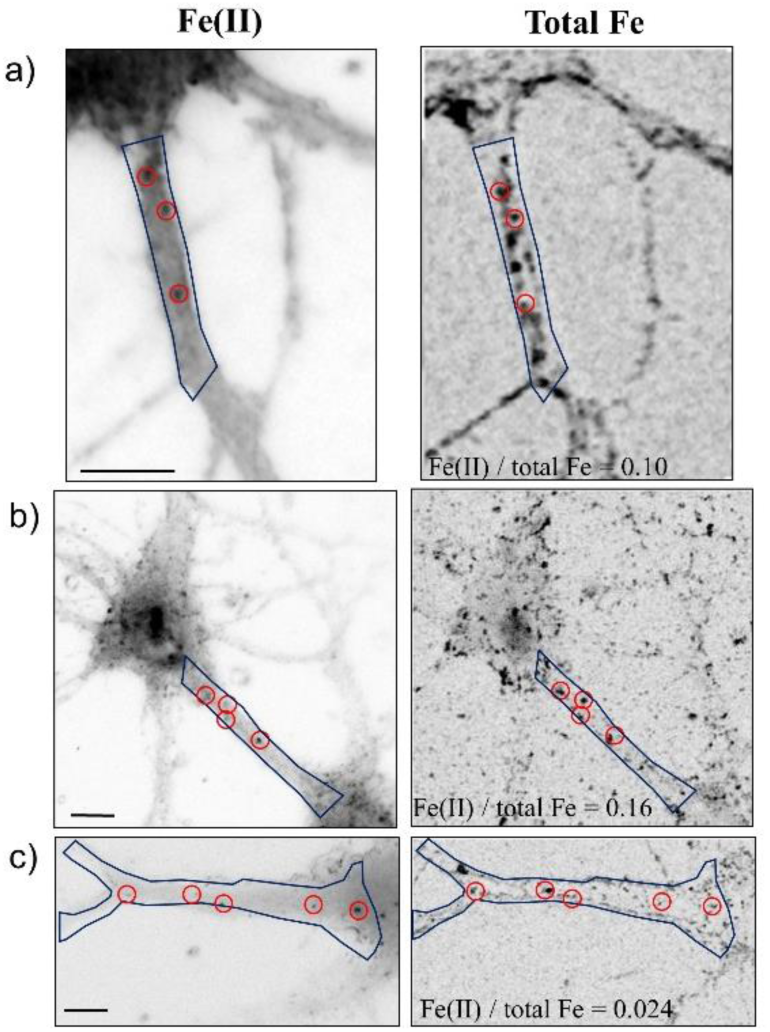
Examples of quantification of labile Fe(II) over total iron content in dendrites after exposure to 100 µM FAS for 1.5 h. The red circle ROIs indicate Fe(II) localizations determined by FerroFarRed™ fluorescence imaging, while the dark blue polygons represents the total selected dendrite areas. In these three examples, the ratio of the sum intensity of the Fe(II) ROIs to the total iron intensity in the selected dendritic areas ranged from 0.024 to 0.16.

Identifying and quantifying the different oxidation states of iron is critical to understanding iron reactivity in cells. Several methods have been developed for this purpose. One very effective technique is X-ray absorption microspectroscopy, which has been widely used to investigate iron speciation in studies of neurological disorders [30]. In most cases, micro-XAS was able to discriminate between different Fe(III) compounds in cells, but not able to detect Fe(II) species. Although XAS micro-spectroscopy is a very powerful method for accurately determining the oxidation states of iron in biological samples, the approach we propose may have some advantages. Fe(II) species are transient in cells and are rapidly oxidized to the less reactive Fe(III) species. Therefore Fe(II) species are easier to observe in living cells than in fixed samples. Moreover, micro-XAS are time consuming experiments, requiring the scanning of the sample at various energies to image the distribution of different chemical states of the elements. Our method, combining confocal microscopy and SXRF, enables to reduce the synchrotron beam time used by performing only a single scan.

Fluorescence imaging methods using fluorescent iron sensors have been developed and extensively used to study the pool of labile iron in cells [8,11,17,18,31]. These approaches are of considerable interest for investigating the transient and biologically reactive pool of labile iron. However, the proportion of Fe(II) labile iron pool to the total iron content cannot be evaluated since the total iron content is not detected. We present in this article the proof of principle for the combined evaluation of Fe(II) and total iron content in cultured cells. Correlative quantitative imaging defines a concentration range for Fe(II) content in dendrites in the 2.4 to 16% range (Figure 5). These values should be considered as maximum Fe(II) values since our method assumes that iron in Fe(II) puncta is only present as Fe(II). The labile Fe(II) over total Fe content varies according to the selected areas, depending on the number of Fe(II) puncta and total iron in the selected area, but overall the method gives a valuable estimation of the proportion of Fe(II) that is present in the lysosomes. As discussed above, the subcellular determination of labile iron content versus total iron content is difficult to assess because the measurements are based on different spectro-imaging methods. However, our results can be compared with the published literature on separate measurements. For example, the total iron concentration in neurons is typically in the range of 500 µM [32] and the kinetically labile iron pool is present in the cytosol at concentrations between 1 and 7 µM [1]. Our quantitative results are consistent with these data, considering that in our experiment neurons were exposed to excess Fe(II).

Labile iron in its ferrous state Fe(II) can participate in Fenton reactions due to its high reactivity, resulting in the generation of ROS. This process can damage cells, and iron redox chemistry has been implicated in the development of neurodegenerative diseases, including Alzheimer’s and Parkinson’s [1,33]. Disturbances in Fe(II) levels can therefore have a significant impact on neurological health. While total iron measurements provide a general overview of iron content, the distinction between Fe(II) and Fe(III) allows a more detailed understanding of iron metabolism and its direct impact on brain function. We have shown that hippocampal neurons can cope with excess labile Fe(II) by lysosomal storage followed by retrograde and anterograde transport along the axon and dendrites. However, it is possible that this process becomes pathological. In particular, excessive lysosomal iron content in neurons can lead to oxidative stress, resulting in neuronal damage and cell death [21]. Previous research has shown that iron accumulates in the hippocampus of AD patients [7], and that lysosomes also accumulate significantly near extracellular amyloid plaques in AD, inhibiting their degradation capacity and disrupting retrograde axonal transport [34]. The presence of high levels of Fe in lysosomal accumulations in AD patients merits further investigation.

## Conclusion

Our data show that after exposure to labile Fe(II), intra-neuronal iron is distributed within somas and neurites suggesting iron uptake via iron transport membrane proteins expressed in the soma but also along the entire neuronal processes. Once incorporated into neurons, iron is mainly present in iron-rich clusters, referred to in this article as micrometric puncta, in the soma and neuronal processes. Most of these iron-rich puncta are composed of non reactive iron, most probably bound to ferritin. A smaller fraction of intracellular iron is present as labile Fe(II) and is located within lysosomes. Lysosomal labile Fe(II) can be transported retrogradely and anterogradely along the axon and dendrites. Although the precise role of labile iron transport in axons and dendrites is not fully known, it is thought to serve several functions, including retrograde transport to the cell soma for detoxification and storage in the form of unreactive iron species, and anterograde transport to meet iron demand at the terminal ends of axons and dendrites. Anterograde transport of Fe can be used for long-distance transport of iron to distant parts of the brain [26]. Finally, we have developed a methodological workflow that allows a straightforward assessment of labile Fe(II) versus total iron content at the subcellular level, using correlative imaging of a fluorescent marker for labile Fe(II) and SXRF imaging of total iron. This method indicates that a small but significant amount of labile Fe(II) is present in dendrites within lysosomes after Fe(II) exposure, ranging from 2.4 to 16% of the total iron content in the samples treated. This method can now be used more generally for *in vitro* studies of iron redox biochemistry.

## Author Contributions

A.K., A.C., C.P. and R.O. designed the experiments. A.K. and A.C. performed the laboratory experiments. A.K., A.C. and A.S. performed synchrotron experiments. A.K. analysed the data and prepared the figures. A.K. and R.O. wrote the first draft of the manuscript. A.K., A.C., A.S., C.P., and R.O. edited the manuscript. All the authors read and approved the final version.

## Acknowledgments

This project was funded by the National Research Council of Thailand through the RGJ-Ph.D. Program (project code: N41A650085). We acknowledge the partenariat Hubert Curien SIAM for funding (project number 50812YC). We would like to thank the Cell Biology Facility from the Interdisciplinary Institute of Neuroscience at the University of Bordeaux for providing the primary rat hippocampal neurons. The fluorescence and confocal microscopy was performed at the Bordeaux Imaging Center, a service unit of the CNRS-INSERM and Bordeaux University, member of the national infrastructure France BioImaging. Synchrotron experiments were performed in the Nanoscopium beamline at the SOLEIL French national synchrotron facility, France.

